# Zebrafish IL-4-like cytokines and IL-10 suppress inflammation but only IL-10 is essential for gill homeostasis

**DOI:** 10.1101/2020.04.09.033837

**Authors:** Federica Bottiglione, Christopher T. Dee, Robert Lea, Leo Zeef, Andrew P. Badrock, Madina Wane, Laurence Bugeon, Margaret J. Dallman, Judith Allen, Adam F. L. Hurlstone

**Affiliations:** School of Biological Sciences, and Lydia Becker Institute of Immunology, Faculty of Biology, Medicine and Health, The University of Manchester, Manchester, UK; Department of Life Sciences, Faculty of Natural Sciences, Imperial College London, London, UK

## Abstract

Healthy fish stocks are central to global food security. Key to fish health is robust immunity at mucosal surfaces, and especially at the gills. However, a balance must be struck between tolerating commensal microorganisms and reacting appropriately toward pathogens. In mammals, IL-4 and IL-13 in concert with IL-10 are essential for balancing immune response to pathogens and suppressing inflammation. Whether their fish counterparts perform similar roles is an open question. Here, we have generated IL-4/13A and IL-4/13B mutant zebrafish and, together with existing IL-10 mutants, characterized the consequences of loss-of-function of these cytokines. We demonstrate that these cytokines are required to suppress inflammation. Further, IL-4/13A and IL-4/13B are required for the maintenance of a Th2-like phenotype in the gills. As in mammals, IL-10 appears to have a more striking anti-inflammatory function than IL-4-like cytokines. Thus, both IL-10 and IL-4/13 paralogues in zebrafish exhibit aspects of conserved function with their mammalian counterparts.

## Introduction

In mammals, interleukin (IL)-4 and −13 are structurally and functionally related cytokines that participate in several physiological processes but are mainly known for stimulating type 2 immune responses (1), characterised by mucus overproduction, IgE antibody production, eosinophilia and differentiation of alternatively activated (M2) macrophages (2). These responses confer protection against parasites but, when inappropriately activated, contribute to the development of asthma and allergic inflammation (1). IL-4 and IL-13 are secreted by CD4^+^ Th2 cells, basophils, eosinophils, mast cells and group 2 innate lymphoid cells (ILC2s) (3). They exert their functions by binding to two types of receptor complexes: a type I receptor, constituted by the IL-4Rα chain and the IL-2R common γ chain (γc) and a type II receptor, which comprises IL-4Rα chain and IL-13Rα1 subunits. Both type I and type II receptors signal through STAT6 transcription factor binding to promoter elements within IL-4/IL-13 responsive genes (4). IL-4 and IL-13 suppress inflammatory responses by antagonising production of TNFα, IL-1β and other pro-inflammatory mediators (5) and act in opposition to IFN-γ, the canonical Th1 (type 1) cytokine (6). Indeed, co-ordinately with inducing type 2 immune response, IL-4 and IL-13 suppress type 1 responses, characterised by immune cell mediated destruction of cells infected with intracellular pathogens (7).

Although IL-4 and IL-13 can negatively regulate inflammatory responses, IL-10 has the more central anti-inflammatory role in mammals, potently suppressing IFN-γ responses. IL-10 can suppress a range of aberrant immune responses including both type 1 and type 2 responses (8). IL-10 is produced by many cell types (9). It regulates CD4^+^ Treg cell differentiation and function, and it is important in maintaining homeostasis at mucosal surfaces (10).

Evidence is emerging that fish immune responses can also be classified as type 1 or type 2 and that T helper cells similar to mammalian counterparts likely do exist (11). Relevant cytokine receptors and downstream signaling pathway molecules are conserved between fish and mammals (12). However, there is still the need to understand how specialized immune cells and cytokines function, in particular in mucosal tissues such as the gills. Fish gills represent an important mucosal surface that is constantly exposed to both waterborne pathogens and commensals and is able to mount immune responses (13,14). Several immune cells including macrophages, neutrophils and lymphocytes have been described in the gills of teleost fish (15). As with all mucosal surfaces, the balance between inflammatory and regulatory responses is critical for the maintenance of tissue homeostasis and barrier integrity.

Two type-2 cytokines that are related to tetrapod IL-4 and IL-13 have been described in teleost fish (16). It is likely that a single *il4/13* gene existed in ancestral jawed vertebrates which has been duplicated in different lineages by whole genome duplication (WGD) and/or tandem duplication events (17). Due to an additional round of WGD event that occurred in fish, teleosts acquired two *il4/13* loci (ohnologues) namely *il4/13a* and *il4/13b* (18). Analysis of these loci shows that teleost *il4/13* genes share important characteristics with genes encoding tetrapod IL-4 and IL-13: their positioning relative to *rad50* and *kif3a* gene neighbors (19); the typical short-chain type 1 cytokine organization as well as conserved structural motifs (18). The identification of putative GATA3 binding motifs in the promoter regions of teleost *il4/13* supports that these genes encode Th2 cytokines (18).

Enriched levels of mRNA encoding IL-4/13A together with the transcription factor GATA3 have been detected in salmonid fish mucosal tissues such as the gills and the skin, indicating these to represent a Th2-skewed environment that protects fish from parasites and inflammatory responses (20). A population of CD4^+^ Th2-like cells and their signature cytokines have also been charaterized in zebrafish gills (21). To date, a comprehensive characterization of fish IL-4/13 paralogues is lacking and whether *il4/13a* and *il4/13b* represent authentic *IL4* and *IL13* genes is still a matter of debate. IL-10 has been described in several fish species (22–25) and its constitutive expression was reported in zebrafish kidney, gut and gills (25), suggesting a role in maintaining homeostasis in these tissues. Exploring the conservation and divergence of IL-4/13 and IL-10 cytokines would further our understanding of fish immune responses and may also provide an alternative model for dissecting aspects of mammalian immunity.

Here we used zebrafish to study the functions of fish IL-4/13A and IL-4/13B cytokines. We generated zebrafish knock-outs for *il4/13a* and *il4/13b* genes and addressed the effects of their loss in both larvae and adult fish immunity. We showed the importance of IL-4/13A and IL-4/13B in suppressing inflammation as well as maintaining a Th2 phenotype in the gills. To gain further insight into the regulation of inflammation, the gills of *il10*-deficient zebrafish were examined, revealing the requirement for IL-10 in maintaining homeostasis in this mucosal tissue.

## Materials and Methods

### Zebrafish care

Zebrafish (*Danio rerio*) were maintained under standard conditions (~28 °C under a 14 h light/10 h dark cycle) within the Biological Services Unit (The University of Manchester) and the Central Biomedical Services (Imperial College London). Embryos were collected and raised in egg water (Instant Ocean salt 60 µg/mL) up to 5 days post-fertilisation (dpf) and then transferred to the main aquarium system. Younger fry were fed powder food and rotifers, while older fry and adults were fed powder food and brine shrimp. All food was supplied by ZM fish food. *il10^e46/e46^* mutant zebrafish were a kind gift from Simon Johnston (University of Sheffield) and were generated by the Sanger Institute through the Zebrafish Mutation Project (26). All regulated procedures received ethical approval from the Institutions’ ethical review boards and were performed under Home Office Licence (project licences PF74F0848 and P5D71E9B0) according to the UK’s Animal Act.

### CRISPR/Cas9-mediated generation of mutant lines

#### Creation of single chimeric guide (g)RNA targeting il4/13a and il4/13b

gRNAs were designed to target the first exon of either *il4/13a* or *il4/13b* zebrafish genes using the Harvard chopchop program (https://chopchop.rc.fas.harvard.edu). The *il4/13a* target site was 5’-GGGTTTTACGTTGAAAGGCA-3’ and *il4/13b* target site was 5’-GAAATCATCCAGAGTGTGAA-3’ and were incorporated into a gRNA template for transcription using PCR. The forward primer contained the target gene specific sequence followed by a constant sequence which overlaps with the remaining sequence of *S. pyogenes* chimeric gRNA DNA template (plasmid #51132; Addgene) while the reverse primer was complementary to the gRNA DNA template (see Table I for primer sequences). PCR amplification of the gRNA DNA template using a high fidelity Phusion Taq polymerase was performed and PCR products were gel purified and used for the synthesis of gRNAs using an Ambion MEGAshortscript T7 kit according to the manufacturer’s instructions.

**Table I.**
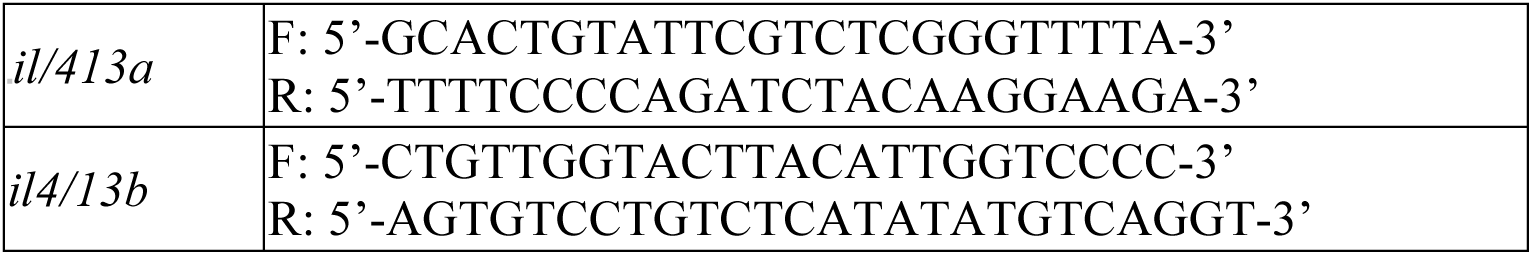
Primer oligonucleotides sequences for PCR

#### Injecting CRISPR reagents

Injection solutions included 30 pg/nL of either gRNA, 250 pg/nL of codon-optimized *S. pyogenes* nls-zCas9-nls mRNA synthesized using a mMESSAGE mMACHINE SP6 kit (Life Technologies) from a pCS2 construct (plasmid #47929; Addgene), 100 pg/nL H2B-mCerulean3 tracer mRNA similarly generated from a pCS2 construct, and 0.05 %(w/v) phenol red to allow visualization of injections. Embryos were injected at the one-cell stage and screened for fluorescence at 24 hpf to identify positively injected fish, which were then raised to adulthood. To identify modified alleles in the F1 progeny for the establishment of *il4/13a* and *il4/13b* zebrafish mutant lines, genomic DNA was amplified by PCR using gene specific primers flanking the targeted site (Table I) and abnormal sized products were gel purified. 1 µL of sample was cloned into a TOPO cloning vector (Invitrogen). Cloning reactions were performed according to the manufacturer’s instructions. Inserts were sequenced by Sanger sequencing using Eurofins Genomics services.

### Whole mount in situ hybridization (WISH)

Three days old embryos were fixed in 4 %(w/v) paraformaldehyde (PFA) in PBS at 4 °C overnight (O/N). Fish were then washed in 0.1% (v/v) Tween20 PBS solution (PBST) and dehydrated through an ascending series of methanol. Digoxigenin (DIG)-labelled RNA antisense probe for zebrafish *mpx* was kindly supplied by Stephen Renshaw (Sheffield) and WISH was performed as described in (27). Embryos fixed in glycerol were mounted on a glass slide and images were collected on a Leica microscope.

### Real-time quantitative (q)PCR

RNA was isolated from homogenised larvae and/or adult tissues using the RNeasy mini kit (Qiagen) and reverse transcribed using the ProtoScript II first strand cDNA synthesis kit (New England Biolabs) with Oligo (dT) primers according to the manufacturer’s instructions. QPCR was performed using SYBR Green JumpStart *Taq* ReadyMix (Sigma), Rox reference dye and cDNA (diluted 1:2). All samples were run in triplicate on 96-well PCR plates (Bio Labs). Negative controls were run without cDNA. Data were analysed by the ΔCt method using *bactin* for normalisation. The primers used for qPCR are listed in Table II.

**Table II.**
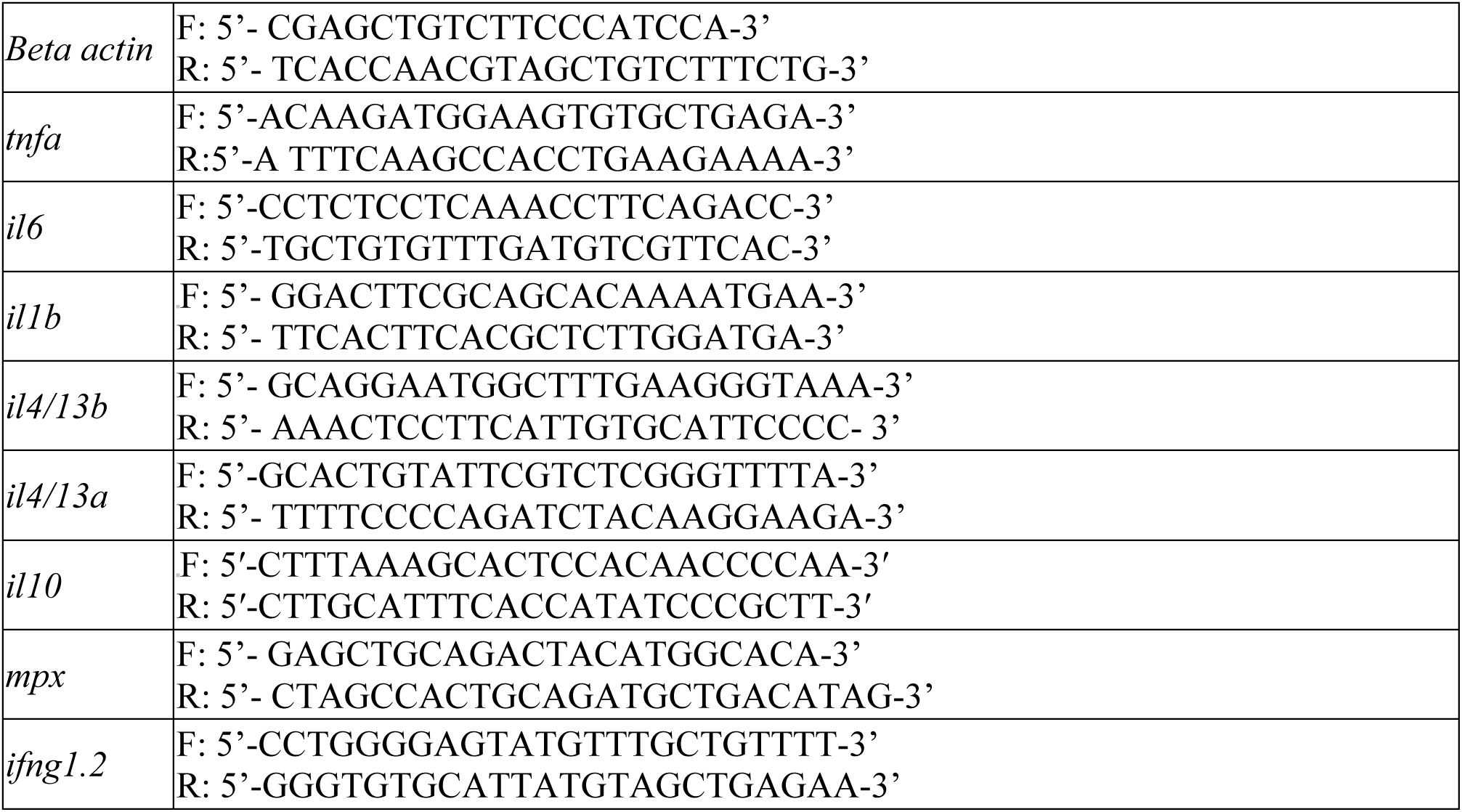
Primer oligonucleotides sequences for q-PCR

### Histology

Whole zebrafish heads were fixed in 4 % (w/v) PFA overnight at 4 °C. Next day the samples were washed in PBS and decalcified in 0.25 M ethylenediaminetetraacetic acid (EDTA) solution for 3 days at room temperature. After decalcification, heads were washed with tap water and placed in 70 % (v/v) EtOH. All specimens were dehydrated and paraffin wax embedded. Sections (5 μm) were cut on a Leica rotary microtome RM2235 and mounted on slides (SUPERFROST PLUS, Menzel-Glaser, Thermo Fisher). Slides were deparaffinised, rehydrated and stained with haematoxylin and eosin. Staining and coverslipping was performed on a Leica workstation operated by the Histology Facility at The University of Manchester. Slides were scanned on a slidescanner (Pannoramic P250 Flash III) and images were processed using 3DHISTECH software.

### RNA sequencing (RNA-seq)

Prior to performing RNA-seq, the integrity of RNA samples was assessed using a 2200 TapeStation (Agilent Technologies) according to the manufacturer’s instructions. Total RNA was submitted to the Genomic Technologies Core Facility (GTCF) of the University of Manchester and libraries were generated using the TruSeq® Stranded mRNA assay (Illumina, Inc.) according to the manufacturer’s protocol. Adapter indices were used to multiplex libraries, which were pooled prior to cluster generation using a cBot instrument. The loaded flow-cell was then paired-end sequenced (76 + 76 cycles, plus indices) on an Illumina HiSeq4000 instrument. Finally, the output data was demultiplexed (allowing one mismatch) and BCL-to-Fastq conversion performed using Illumina’s bcl2fastq software, version 2.17.1.14. Unmapped paired-end sequences were tested by FastQC (http://www.bioinformatics.babraham.ac.uk/projects/fastqc/). Sequence adapters were removed and reads were quality trimmed using Trimmomatic_0.36 (28). The reads were mapped against the reference zebrafish genome (danRer10) and counts per gene were calculated using annotation from ENSEMBL (http://ftp.ensembl.org/pub/current_gtf/danio_rerio/Danio_rerio.GRCz10.84.gtf.gz) using STAR_2.5.3 (29). Normalisation, Principal Components Analysis and differential expression were identified using DESeq2_1.16.1 (30) as genes having a corrected *p* value <0.05 and log2 fold-change >|1|. The RNA-seq datasets reported in this paper have been deposited in the ArrayExpress database (accession number E-MTAB-8958).

### Cluster analysis and functional analysis of differentially expressed genes

Clustering analysis was performed using k-means clustering (Manhattan distance) using the maxdView software. Each cluster was then hierarchically clustered. Clustering was performed on the means of each sample group (log 2) that had been z-transformed (for each gene the mean set to zero, standard deviation to 1). Gene ontology analysis and pathway analysis were performed with Enrich (31) using human orthologues of zebrafish genes obtained from Ensembl genome browser version 99. Gene set enrichment analysis was performed using GSEA software (32). Several gene sets consisting of different immune cell markers, spanning innate and adaptive immune cell types, were generated using the human immunology panel from Nanostring. Normalised enrichment scores (NES) were used to generate heatmaps with Morpheus software (https://software.broadinstitute.org/morpheus). xCell analysis was used to perform cell type enrichment analysis from gene expression data for 64 immune cell types (https://xcell.ucsf.edu).

### Resiquimod (R848) gill challenge

Resiquimod gill challenge was performed as previously described (33) using 5 μL of Resiquimod (0.5 mg/mL; Invivogen) applied to the gills for 5 min. Following completion of the procedure, fish were returned to fresh system water and monitored for their recovery. Fish were then sacrificed with an overdose of MS222 (800 mg/L) at the indicated time point to harvest the gills for RNA extraction.

### Statistical analysis

Statistical analysis was performed using GraphPad Prism version 8 (GraphPad Software Inc.). P-values of less than 0.05 (where necessary corrected for multiple testing) were considered statistically significant. Throughout, error bars represent SEM.

## Results

### Generation of *il4/13a* and *il4/13b* zebrafish mutants

In zebrafish, two similar *il4*-like genes, namely *il4/13a* and *il4/13b*, have been described (18). To generate insight into the function of these two genes, we created mutant lines lacking functional alleles. Specific guide (g)RNAs were designed to target the first exon of either the *il4/13a* or *il4/13b* gene. Each gRNA was co-injected with Cas9-encoding mRNA into one-cell stage zebrafish embryos. In order to identify specific mutations, the gRNA target sites were amplified from F1 offspring and DNA sequencing of cloned PCR products was performed. This revealed an *il4/13a* allele carrying a 28 base pair frame shift mutation (32bp insertion and 4bp deletion) and an *il4/13b* allele carrying a 7 base pair deletion (Figure 1A). Fish carrying the identified mutations were out-crossed and the F2 progeny were genotyped for the presence of mutations. *Il4/13a*^+/−^ and *il4/13b*^+/−^ heterozygous fish were selected and raised to adulthood. In-crosses of heterozygotes were performed to generate homozygous mutants for both *il4/13a* (*il4/13a*^−/−^) and *il4/13b* (*il4/13b*^−/−^) in the F3 generation. Homozygous mutants were found to be viable and they were further in-crossed to maintain mutant lines. In order to generate *il4/13a;b*^−/−^ double mutants, *il4/13a*^−/−^ and *il4/13b*^−/−^ homozygous fish were crossed and their progeny in-crossed. These were also found to be viable and they were further in-crossed to maintain the line.

**Figure 1.**
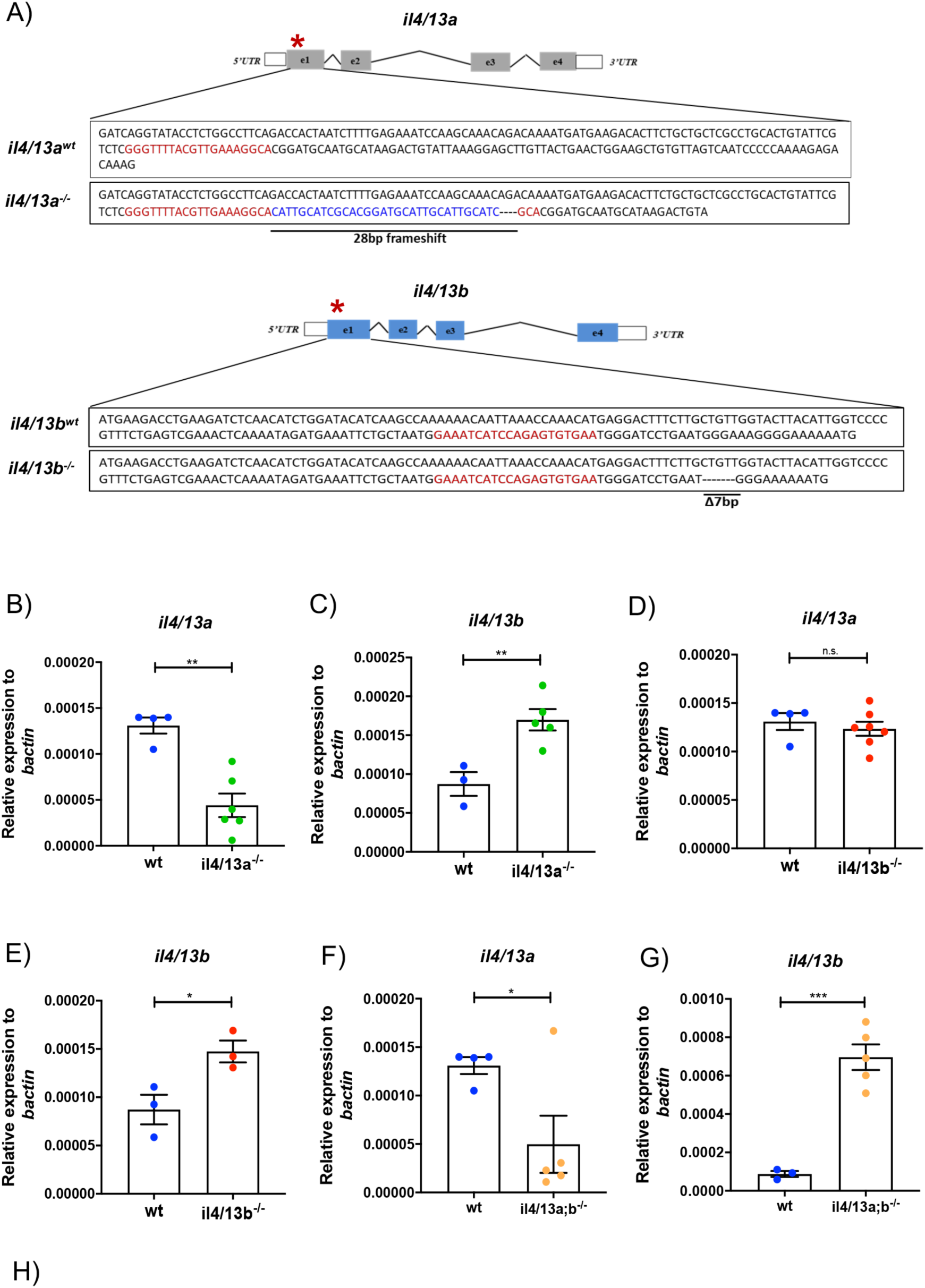
Generation of *il4/13a^−/−^* and *il4/13b^−/−^* zebrafish mutants by CRISPR/Cas9. (A) Zebrafish *il4/13a* and *il4/13b* genes. (B)-(G) qPCR analysis showing the levels of *il4/13a* and *il4/13b* transcripts in wildtype, *il4/13a^−/−^, il4/13b^−/−^* and *il4/13a;b*^−/−^ 5dpf larvae. Gene expression was normalized to the expression of βactin. Error bars represent SEM; n>3, *p<0.05, **p<0.01, ***p<0.001. (H) Sequence of wildtype and mutated IL-4/13A and IL-4/13B proteins. Underlined is the unchanged portion of the protein sequence in mutant animals. Yellow-shaded ‘R’ indicates the arginine residue. The asterisks indicate a stop codon.

To evaluate whether the mutations affected cognate mRNA levels, qPCR analysis was performed. Significant reduction of *il4/13a* mRNA in *il4/13a^−/−^* embryos was observed (Figure 1B) suggesting mRNA degradation by the nonsense-mediated decay (NMD) pathway (34). Perhaps as a consequence of NMD, an increase of *il4/13b* mRNA (1.6 fold) was detected in *il4/13a*^−/−^ embryos (Figure 1C), suggesting a potential mechanism of genetic compensation as recently described (35). No changes in the levels of *il4/13a* mRNA were observed in *il4/13b^−/−^* larvae (Figure 1D). The expression of *il4/13b* mRNA was found increased in *il4/13b^−/−^* larvae (Figure 1E), suggesting that the effects of the mutation might only be evident at the translational level and that increased mRNA might be an attempt to compensate for loss-of-function of the IL-4/13B protein. qPCR analysis also revealed that the expression of *il4/13a* was significantly reduced in *il4/13a;b*^−/−^ embryos (Figure 1F) whereas the expression of *il4/13b* was significantly increased (Figure 1G), in agreement with what was observed in the single mutants.

The identified mutations were predicted to introduce a premature termination codon (PTC) in both *il4/13a* and *il4/13b* genes leading to the formation of severely truncated proteins. Indeed, only 16 and 28 amino acids remained in the truncated IL-4/13A and IL-4/13B proteins, respectively (Figure 1H). Previous analysis of zebrafish protein sequences revealed conservation of an arginine in the αC helix (36) which is critical for the binding of human IL-4 to the IL-4Rα receptor. The mutations identified in IL-4/13A and IL-4/13B proteins abolished this residue implying that the proteins should not be able to bind to their receptors and thus lack any functional activity.

### Disruption of both il4/13a and il4/13b genes leads to a pro-inflammatory phenotype in larvae

The functions of *il4/13a* and *il4/13b* genes were first investigated in larvae, which possess an innate immune system only (37). While a modest increase of *tnfα*, *il6* and *ifng1-2* mRNA encoding pro-inflammatory cytokines was observed in *il4/13a*^−/−^ and *il4/13b*^−/−^ single mutants, more pronounced up-regulation was observed in *il4/13a;b*^−/−^ double mutants (Figure 2A), indicating that *il4/13a* and *il4/13b* genes have a fundamental albeit redundant role in suppressing inflammation.

**Figure 2.**
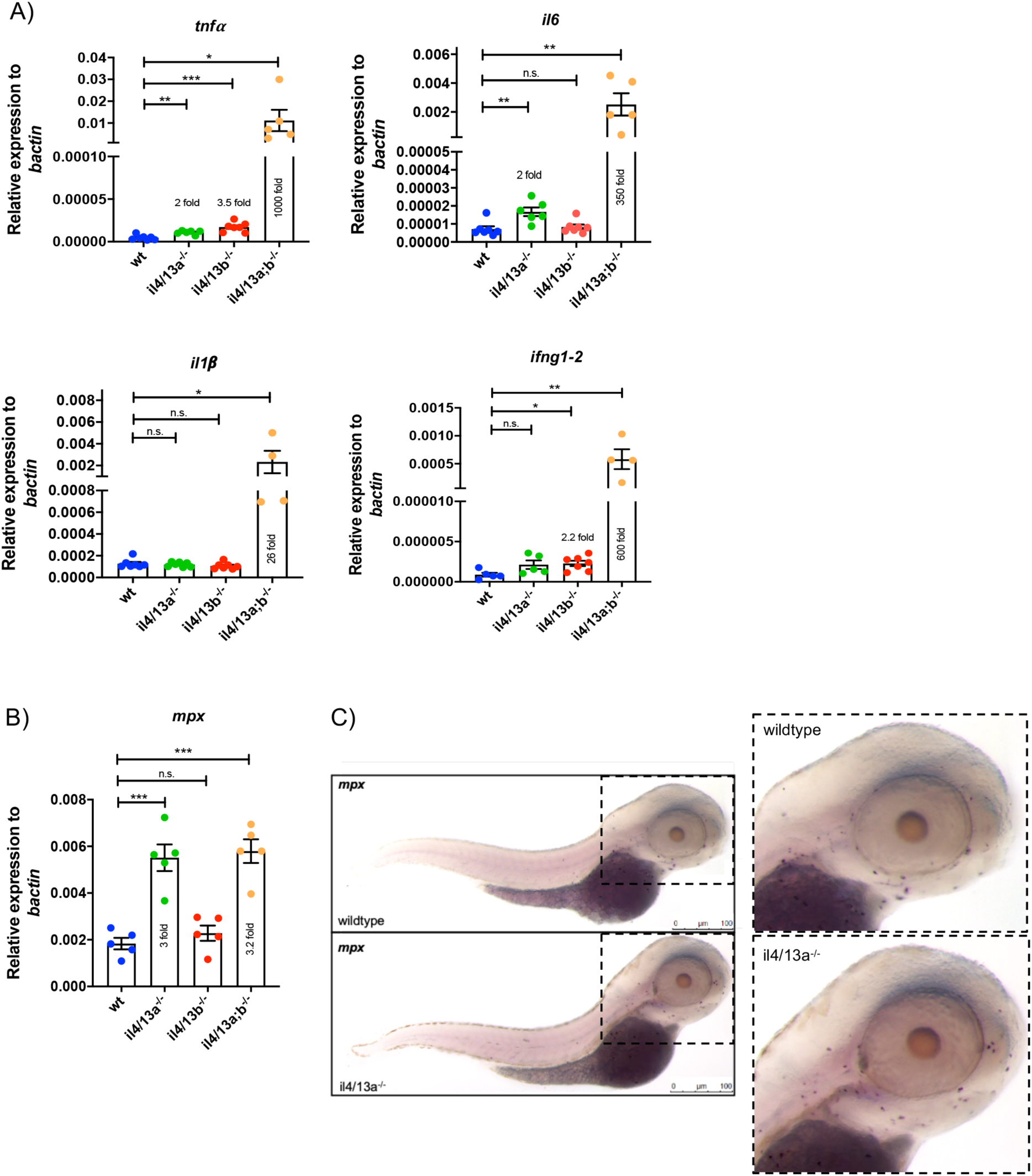
Increased levels of pro-inflammatory cytokines in *il4/13a;b^−/−^* larvae. (A) and (B) qPCR analysis showing levels of *il1β*, *il6*, *tnfα*, *ifng1-2* and *mpx* mRNA in wildtype, *il4/13a^−/−^, il4/13b^−/−^* and *il4/13a;b^−/−^* 5dpf larvae. Dots indicate individual fish. Gene expression was normalized to the expression of βactin. Error bars represent SEM; n>3, *p <0.05, **p<0.01, ***p<0.001. (C) Representative image of *mpx in situ* hybridization in wildtype and *il4/13a^−/−^* 3dpf larvae (n=12-15). Purple dots indicate *mpx+* cells. Scale bar 100µm.

Next, we investigated whether the inflammatory phenotype was accompanied by an increase of neutrophils by measuring the levels of myeloperoxidase *mpx* transcript in larvae. A significant upregulation of *mpx* was detected in *il4/13a;b*^−/−^ double mutants (3.2 fold) supporting the inflammatory phenotype. A similar increase of *mpx* was observed in *il4/13a*^−/−^ larvae (3 fold) but not in *il4/13b*^−/−^ larvae (Figure 2B), suggesting the increase in *mpx* transcript might be due to loss of functional IL4/13A. Thus, in situ hybridization for *mpx* mRNA was performed on wildtype and *il4/13a*^−/−^ larvae to further investigate changes in neutrophils. However, no difference in neutrophil numbers or distribution were detected in *il4/13a*^−/−^ embryos (Figure 2C).

### Transcriptome analysis of the gills reveals a shift towards type 1 immunity in *il4/13a;b^−/−^* mutants

Our previous work demonstrated the expression of *il4/13a* in the gill tissue of adult zebrafish as well as *il4/13b* expression in gill-resident CD4^+^ T cells, suggesting the presence of a Th2-skewed environment in the gills (21). In order to investigate the gene expression profile of gills from *il4/13a^−/−^*, *il4/13b^−/−^*and *il4/13a;b^−/−^* mutant animals, adult gill tissue was harvested, the RNA isolated and RNA sequencing (RNA-seq) undertaken. Principal component analysis (PCA) revealed clear segregation and clustering of mRNA profiles according to genotype (Supplemental Figure 1A). 5,619 differentially expressed genes were identified in gills of mutant animals compared to wildtype (Supplemental Table 1). In order to identify coordinated changes in gene expression within our samples, hierarchical clustering was performed. Differentially expressed genes were grouped in ten clusters that were subjected to gene ontology analysis using Enrichr (31), inputting human orthologues of zebrafish genes. Enriched gene ontologies were mainly associated with immune related biological processes (Supplemental Figure 1B).

In order to characterize the transcriptional profile of *il4/13a;b*^−/−^ double mutants, we focused on the genes belonging to cluster 2 that were more highly expressed in *il4/13a;b*^−/−^ double mutants. Gene ontology analysis revealed that these genes were mainly associated with negative regulation of type 2 immune responses, Th1 differentiation, and interferon gamma (IFN-γ) signaling (Figure 3A). Gene set enrichment analysis (GSEA) corroborated the inference that enhanced Th1 differentiation, diminished Th2 differentiation and increased IFN-γ signaling was occurring in the gills of *il4/13a;b*^−/−^ double mutants (Figure 3B). Moreover, qPCR confirmed increased levels of *ifng1-2* mRNA in the gills of *il4/13a;b*^−/−^ double mutants (Figure 3C). All together these data suggest that in absence of both *il4/13a* and *il4/13b* there might be a shift towards type-1 immunity, comparable to the role of IL-4 in mammals.

**Figure 3.**
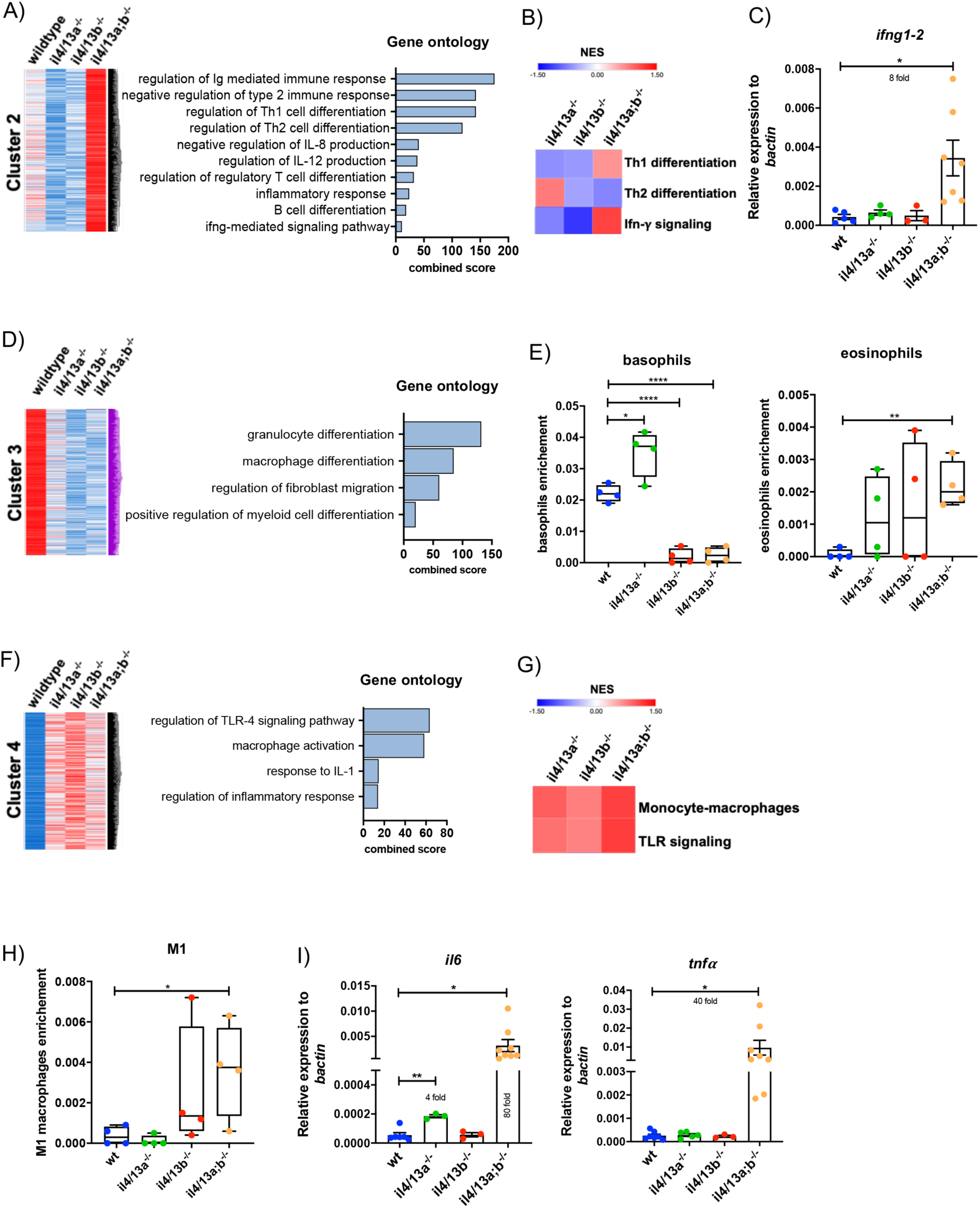
Transcriptome analysis of *il4/13a^−/−^, il4/13b^−/−^* and *il4/13a;b^−/−^* gills. (A), (D) and (F) Clustering of differentially expressed genes identified in gills harvested from 6 month old fish. Blue indicates low expression, red indicates high expression and white indicates unchanged expression. Bar graphs show significantly enriched ontologies for each cluster of genes. (B) and (G) Heatmap of normalized enrichment scores (NES) for the indicated gene sets generated by gene set enrichment analysis (GSEA) in mutant gills compared to wildtype gills. (C) and (I) qPCR analysis showing transcript levels of *ifng1-2*, *il6* and *tnfa* in wildtype and mutants gills. Gene expression was normalized to the expression of βactin. Error bars represent SEM; n>3, *p <0.05, **p<0.01, ***p<0.001, ****p<0.0001. (E) and (H) Enrichment score of basophils, eosinophils and M1 macrophages obtained by xCell analysis.

Next, we sought to investigate the gene signature of cluster 3 as these genes were down-regulated in gills from *il4/13a^−/−^*, *il4/13b^−/−^* and *il4/13a;b^−/−^* animals. Gene ontology analysis revealed that genes from cluster 3 were mainly associated with differentiation of myeloid cells as well as regulation of fibroblast migration (Figure 3D). To characterize potential enrichment of immune cell populations, xCell analysis was performed. Among myeloid-derived cells, basophils were predicted to be significantly reduced in gills from *il4/13b*^−/−^ single mutant and *il4/13a;b*^−/−^ double mutants, whereas an increase of eosinophils was predicted for *il4/13a;b*^−/−^ double mutants at least (Figure 3E).

Genes belonging to cluster 4 were found highly up-regulated in gills from *il4/13a*^−/−^, *il4/13b*^−/−^ and *il4/13a;b*^−/−^ animals and they were found associated with processes such as toll-like receptor 4 signaling, macrophage activation, response to interleukin-1 as well as regulation of inflammatory response (Figure 3F). Accordingly, an enrichment of monocytes-macrophage expression profile as well as TLR signaling signature was detected by GSEA (Figure 3G). Moreover, xCell analysis predicted an increase of M1 macrophages in the gills of *il4/13a;b*^−/−^ double mutants (Figure 3H). To further validate these data we measured expression of pro-inflammatory cytokines in the gills by qPCR and found increased expression of *il6* (80 fold) and *tnfα* (40 fold) mRNA in the gills of *il4/13a;b*^−/−^ double mutants (Figure 3I), suggesting overall heightened inflammatory activity.

### Loss of *il4/13a* and *il4/13b* does not affect gill morphology whereas lack of functional *il10* does

To further evaluate the effects of *il4/13a* and *il4/13b* on zebrafish gill homeostasis, histology was performed. No significant morphological changes were observed in the gills of *il4/13a^−/−^*, *il4/13b^−/−^* or *il4/13a;b^−/−^* fish compared to wildtype (Figure 4A), suggesting that loss of functional IL-4/13A and IL-4/13B does not affect tissue morphology despite a shift toward type 1 immunity. For comparison, we examined the gills of *il10^e46/e46^* mutants lacking functional IL-10, which have also been reported to shift towards type 1 immune responses (38). A thickening of the gill arch and filaments, as well as lamellar epithelial changes were detected in the absence of functional IL-10, indicating a role for this cytokine in maintaining gill homeostasis in adult zebrafish.

**Figure 4.**
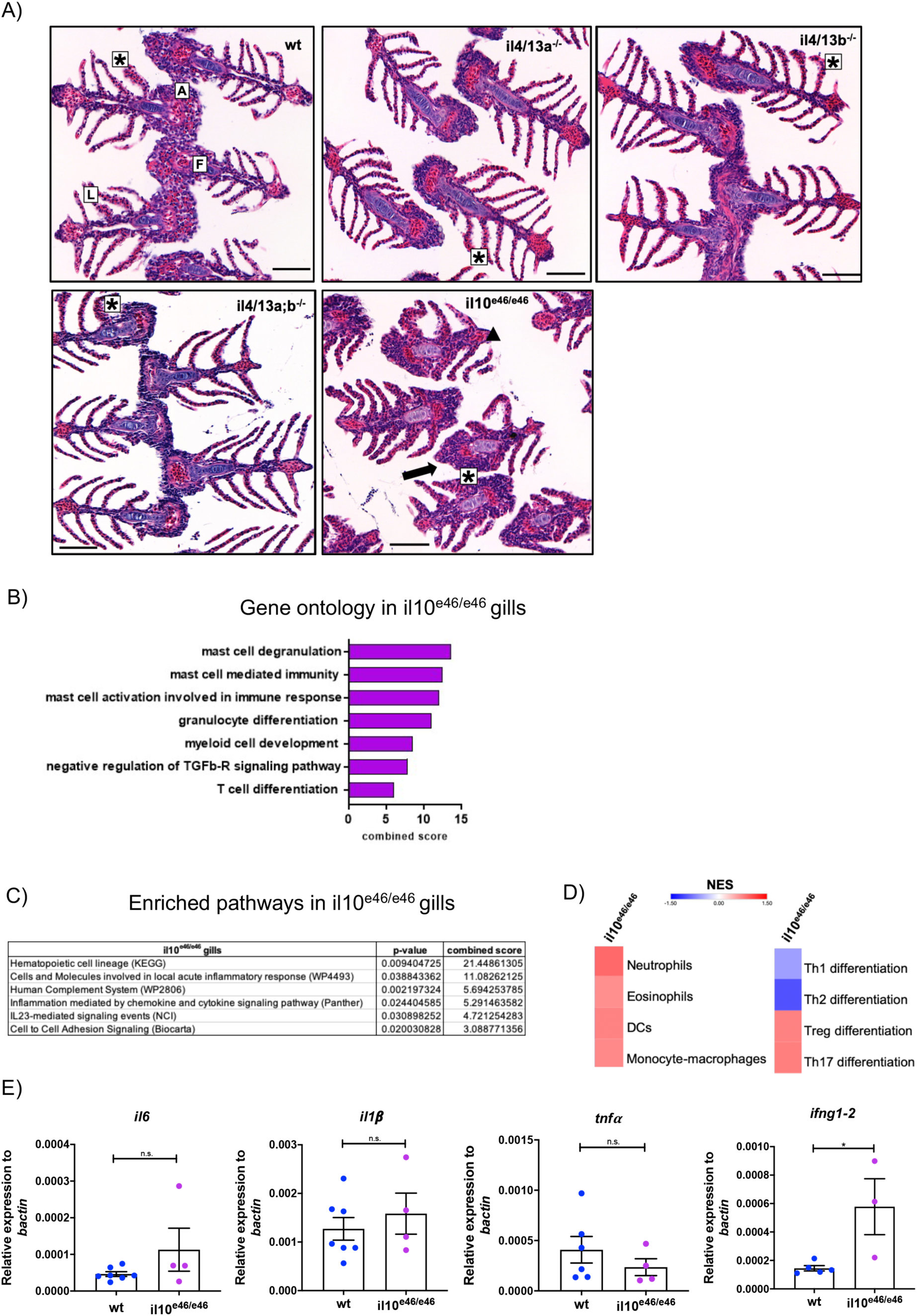
Mutation of il10 induces morphological changes in the gills of adult zebrafish. (A) Representative images of sections of gills from wildtype, *il4/13a*^−/−^, *il4/13b*^−/−^, *il4/13a;b*^−/−^ and *il10*^e46/e46^ animals stained with hematoxylin and eosin. Gill arch (a), filaments (f) and lamellae (l) are indicated. Lamellar epithelial cell alteration (asterisks), filament thickening (triangle) and arch thickening (arrow) are indicated. (B) Bar graph showing significantly enriched ontologies in gills of *il10*^e46/e46^ animals. (C) Table showing significantly enriched pathways in gills of *il10*^e46/e46^. Combined scores and p-values are indicated. (D) Heatmap of NES for indicated gene sets generated by GSEA in gills of *il10e46/e46* compared to wildtype animals. (E) qPCR analysis showing transcript levels of *il1β*, *il6, tnfα* and *ifng1-2* in gills of *il10*^e46/e46^ compared to wildtype animals. Gene expression was normalized to the expression of βactin. Error bars represent SEM; n>3, *p <0.05.

To further investigate the role of *il10* in gill homeostasis, RNA-seq was performed on RNA extracted from the gills of *il10*^*e46/e46*^ mutant animals. Again, PCA confirmed a distinct expression profile for *il10*^*e46/e46*^ mutant animals (Supplemental Figure 1A). Gene ontology analysis was performed using Enrichr and an enrichment of immune-related ontologies, particularly related to mast cell and granulocyte differentiation, was observed (Figure 4B). We also performed pathway analysis using Enrichr and found further evidence for an ongoing inflammatory response, highlighting the action of the complement system, chemokine and cytokine signaling as well as other mediators of inflammation (Figure 4C). In order to investigate potential enrichment by immune cell populations, GSEA was performed. Enrichment of neutrophils, eosinophils, dendritic cells and monocyte-macrophages as well as TLR signaling was predicted in gills from *il10*^*e46/e46*^ mutant animals. Moreover, a depletion of Th1 and Th2 cells but an enrichment of Treg and Th17 was also predicted (Figure 4D). In order to validate these data, we measured gene expression changes by qPCR analysis. No significant changes were observed in the levels of *il1β*, *tnfα*, and *il6* mRNA encoding inflammatory cytokines but an increase of *ifng1-2* mRNA was detected in *il10*^*e46/e46*^ gills compared to wildtype (Figure 4E).

### Loss of *il4/13a* and *il4/13b* leads to an enhanced type-1 response to Resiquimod in the gills of adult zebrafish

We next sought to investigate the importance of *il4/13a* and *il4/13b* in modulating an inflammatory response in the gills to an exogenous source of irritant. In order to induce gill inflammation, we used an intervention we have recently established, whereby a solution of Resiquimod (R848), a synthetic compound that mimics viral ssRNA and interacts with TLR7/8 in mammals (39), is applied directly to the gills (33). Gene expression was analyzed by qPCR at 1 h and 8 h in order to evaluate the kinetics of the response (Figure 5A). An inflammatory response was observed in the gills of wildtype, *il4/13a^−/−^* and *il4/13b^−/−^* animals after 1 h as shown by significantly enhanced levels of *il1β, il6, tnfα* and *ifng1-2* mRNA encoding inflammatory cytokines in R848 treated fish compared to untreated controls (Figure 5B). However, no significant difference was observed between mutants and wildtype. Next, we investigated the later inflammatory response (8 hours post-treatment). We observed a comparable down-regulation of mRNA for the inflammatory cytokines *il1β, il6* and *tnfα* in both wildtype and mutant fish compared to the 1 h time point. *ifng1-2* displayed a different kinetics being significantly up-regulated in both wildtype and mutant fish still at 8 h (Figure 5B). Again, this was comparable for wildtype and mutant fish. All together these data suggest that neither loss of *il4/13a* nor loss of *il4/13b* impairs the response to R848-induced inflammation in the gills.

**Figure 5.**
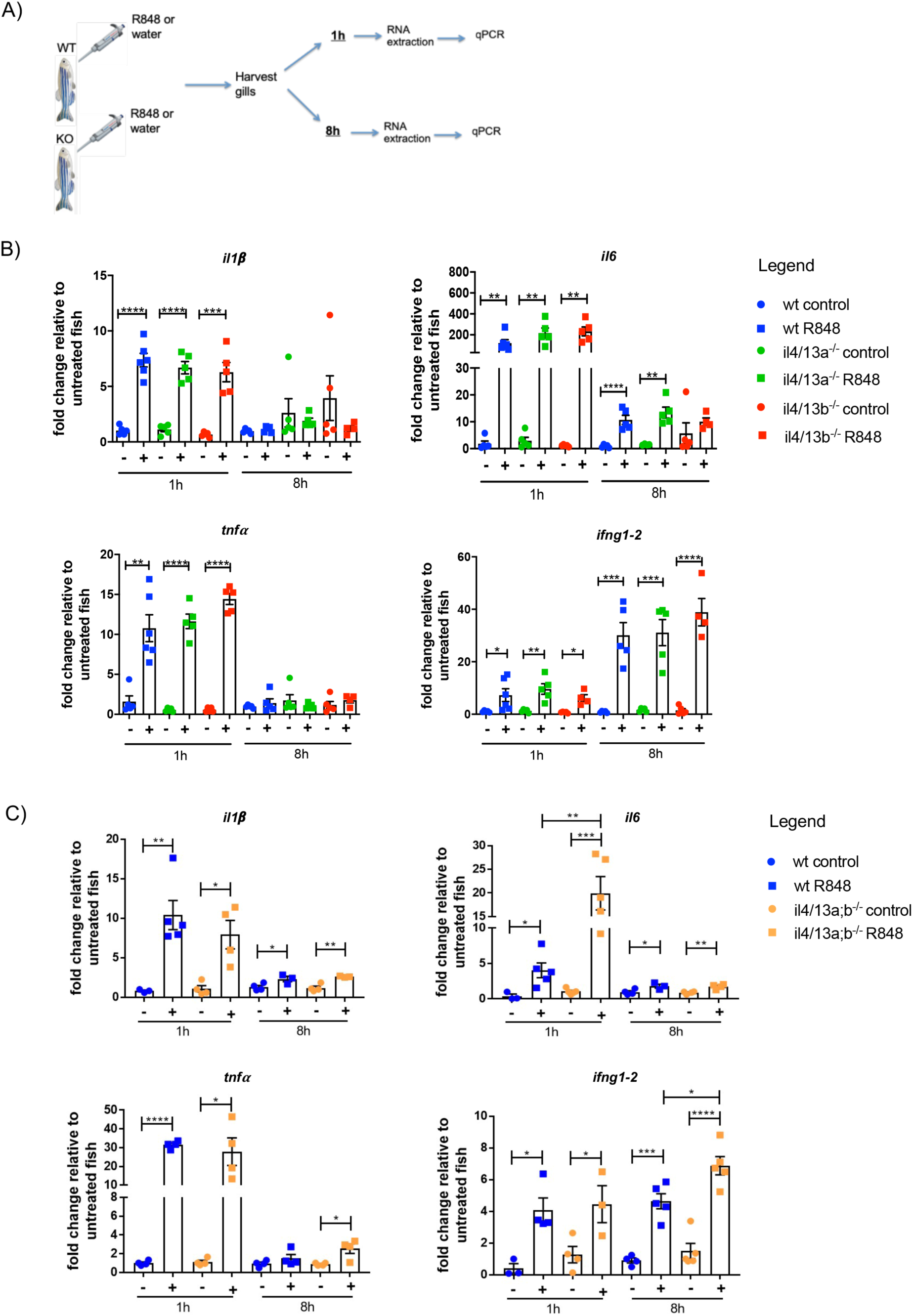
Resiquimod induced an enhanced type-1 response in the gills of *il4/13a;b^−/−^* double mutants. (A) Schematic overview of Resiquimod (R848) gill challenge. (B) and (C) qPCR analysis showing transcript levels of *il1β*, *il6, tnfα* and *ifng1-2* in gills from wildtype, *il4/13a^−/−^, il4/13b^−/−^* and *il4/13a;b^−/−^* animals following R848 challenge for 1 and 8 hours. Gene expression was normalized to the expression of βactin. Fold change is shown. Error bars represent SEM; n>3, *p <0.05, **p<0.01, ***p<0.001, ****p<0.0001.

Next, we evaluated the response to R848 in the absence of both *il4/13a* and *il4/13b*. Resiquimod challenge was performed on the gills of *il4/13a;b^−/−^* double mutants for 1 h and 8 h. Significantly enhanced levels of *il1β*, *il6, tnfα* and *ifng1-2* mRNA were observed in the gills of wildtype and *il4/13a;b^−/−^* animals after 1 h. Moreover, levels of *il6* were significantly higher in *il4/13a;b^−/−^* compared to wildtype (Figure 5C). At 8 h post-treatment, the levels of *il1β*, *il6* and *tnfα* were reduced almost to base line in both wildtype and double mutants. However, *ifng1-2* levels were still elevated and again significantly higher in the gills of *il4/13a;b^−/−^* double mutants compared to wildtype (Figure 5C), suggesting an enhanced type-1 response in agreement with the phenotype observed in the steady state.

### Disruption of *il10* leads to an enhanced inflammatory response to Resiquimod in the gills of *il10^e46/e46^* adult zebrafish

Loss of *il10* resulted in marked alteration of gill morphology in the steady-state and changes in gene expression indicative of smoldering inflammation. We predicted that inflammatory responses to exogenous irritants would be exaggerated in the absence of *il10*. Therefore, the gills of *il10*^*e46/e46*^ fish were challenged with R848 for 1 h and 8 h as described before. Significant upregulation of *il1β, il6, tnfα* and *ifng1-2* mRNA were observed in the gills of stimulated wildtype and *il10*^*e46/e46*^ fish compared to untreated controls after 1 h. However, no difference was observed in the response between wildtype and mutant fish (Figure 6A), suggesting that loss of *il10* does not affect the early phase of an inflammatory response. Next, we evaluated the later response (8 h post-treatment) which revealed that compared to wildtype, *il1β, il6* and *tnfα* transcripts were still significantly upregulated in gills from *il10*^*e46/e46*^ animals and *ifng1-2* transcript levels were even more pronounced (Figure 6A), indicating a prolonged inflammatory response in the absence of *il10*. In keeping with the exaggerated inflammatory response at 8 h post-treatment, thickening of gill filaments and lamellae were observed in gills from R848 treated *il10*^*e46/e46*^ animals but not wildtype animals (Figure 6B). All together these data suggest that loss of *il10* gene induced an exaggerated inflammatory response in the gills following R848 stimulation.

**Figure 6.**
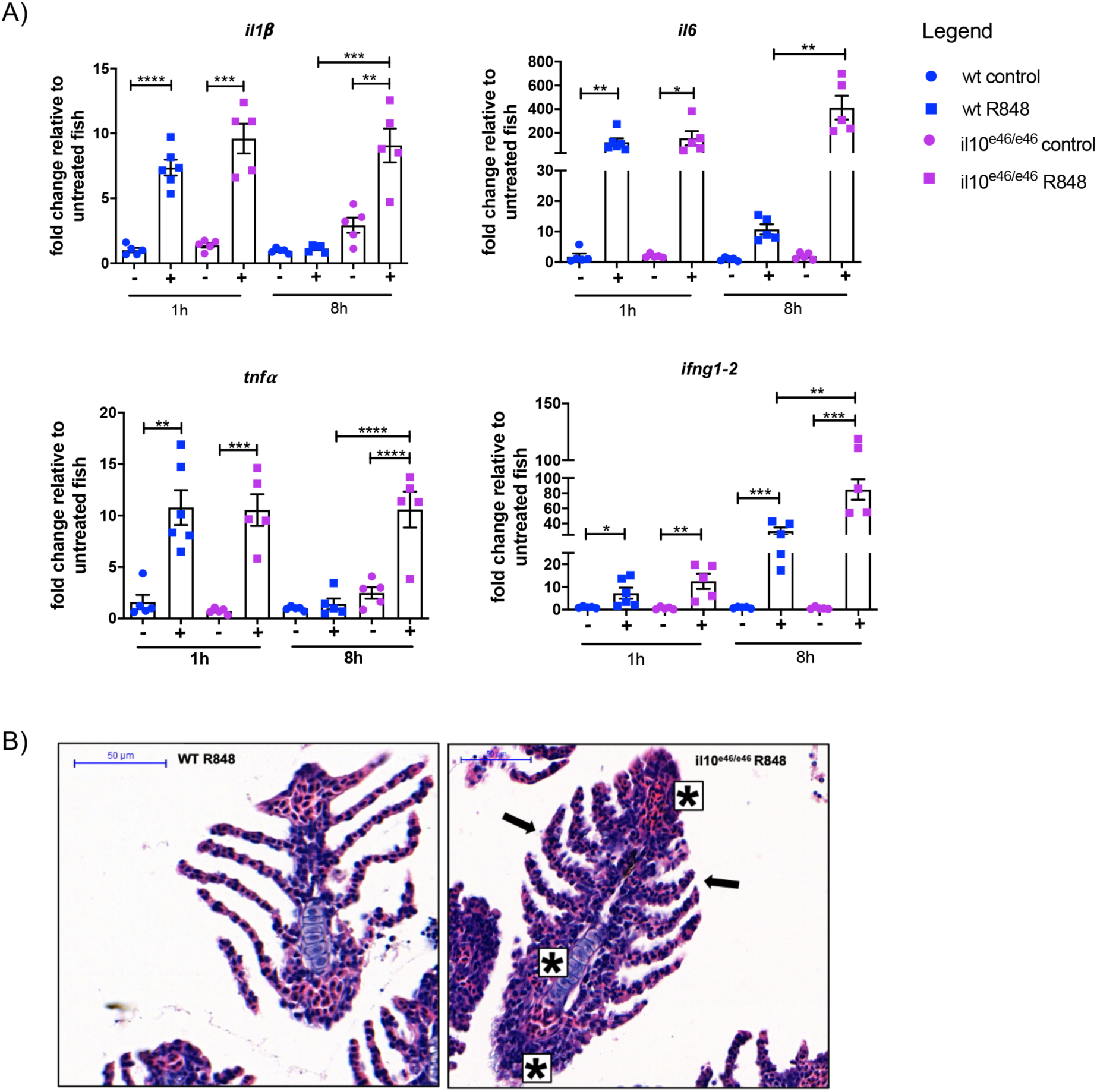
Prolonged inflammatory response in *il10*^*e46/e46*^ gills following Resiquimod stimulation. (A) qPCR analysis showing transcript levels of *il1β*, *il6, tnfα* and *ifng1-2* in gills from wildtype and *il10*^*e46/e46*^ animals following R848 challenge for 1 and 8 hours. Gene expression was normalized to the expression of βactin. Fold change is shown. Error bars represent SEM; n>3, *p <0.05, **p<0.01, ***p<0.001, ****p<0.0001. (B) Representative images of sections of gills from wildtype and *il10*^*e46/e46*^ animals stained with hematoxylin and eosin following 8 hours R848 challenge. Epithelial cell alteration in the lamellae (black arrows) and gill filament (stars) are indicated in gills from *il10*^*e46/e46*^ animals. Scale bar, 50µm.

## Discussion

Fish are central to global food security and advances in fish immunology would help improving fish health and preventing diseases, having a positive impact on aquaculture. Considerable progress has been made in recent years with many genes and cell components of both innate and adaptive immune system being characterized in fish (11,40,41). However, our understanding of immune responses, especially in the gills, where health issues greatly impact farmed fish production (42,43), lacks molecular precision that could help the development of effective vaccines and prophylaxis.

In this study, we have used the zebrafish as a model organism to shed light on fish *il4/13a* and *il4/13b* and characterized the consequences of loss-of-function of these genes under both homeostatic and inflammatory conditions. Fish lacking *il4/13a* and *il4/13b* were viable, fertile and did not show any overt phenotype, indicating that such cytokines are not essential for survival and development in zebrafish. Similar findings were observed in IL-4^−/−^ and IL-13^−/−^ mice that were healthy and displayed no phenotypic abnormalities (44).

We found that disruption of both *il4/13a* and *il4/13b* induced a pro-inflammatory state indicating the role for IL-4/13A and IL-4/13B in suppressing inflammation in zebrafish larvae. An anti-inflammatory role for fish type-2 cytokines has been previously demonstrated in trout where the treatment of head kidney cells with recombinant IL-4/13A and IL-4/13B2 lead to the down-regulation of proinflammatory cytokines (16), resembling the situation in mammals (5). Neutrophils are one of the main components of the zebrafish innate immune system and the first cells to respond to inflammation (45). We found significantly elevated levels of the neutrophil marker *mpx* in *il4/13a;b^−/−^* double mutants supporting the inflammatory phenotype. The increase in *mpx* transcripts in the *il4/13a^−/−^* single mutants indicates that *il4/13a* may have a potential neutrophil-specific effect, albeit one that does not drive a corresponding change in neutrophil number. Effects on *mpx* transcript were not observed in the *il4/13b^−/−^* mutant, suggesting a divergent function for these paralogues in neutrophils in zebrafish. In mammals, neutrophils express IL-4 type I receptor (46,47) and recent evidence suggests that IL-4 and IL-13-mediated signaling in both mouse and human neutrophils inhibits their migration and effector functions (48,49), in order to reduce inflammation and tissue damage during type-2 responses (50). To date, there is no definitive evidence for a role for fish *il4/13* paralogues in controlling neutrophils. The generation of an *il4/13a;b^−/−^;mpx:EGFP* zebrafish line would be valuable to address neutrophil behavior in the absence of functional IL-4/13A and IL-4/13B.

The zebrafish gill mucosa has been described as a source of a resident population of CD4-1+ Th2-like cells, enriched in *gata3* and *il4/13b* expression (21). In order to shed light on the function of *il4/13a* and *il4/13b* in zebrafish gills, transcriptome analysis was performed. We found an enrichment of genes implicated in Th1 differentiation and IFN-γ signaling in the gills of *il4/13a;b*^−/−^ double mutants, further supported by qPCR analysis. These data suggest that *il4/13a* and *il4/13b* are required for the maintenance of a Th2-like phenotype in the zebrafish gills, and their loss leads to an imbalance towards a type-1 immunity. In support of our findings, evidence for a type-2 immunity was previously reported in fish. In zebrafish, interaction of IL-4/13A with IL-4Rα on B cell surfaces promote B cell proliferation and antibody production (51); in trout, treatment of head kidney cells with recombinant IL-4/13 proteins downregulates IFN-γ (16) and an upregulation of *il4/13* paralogues was observed in the gills of Atlantic salmon following parasitic infection (52).

The transcriptome data predicted a reduction of basophils and increase of eosinophils in the gills of *il4/13a;b^−/−^* double mutants. In mammals, basophils and eosinophils represent innate sources of IL-4 and IL-13 and their infiltration into tissues is one of the hallmarks of type-2 immune responses (53). Basophils have been described in teleosts (54) and a conserved role for eosinophils was found in response to helminth antigens and infection (55). It might be speculated that decreased basophils and increased eosinophils in the gills of *il4/13a;b^−/−^* double mutants is somehow connected, perhaps reflecting a divergent effect of IL-4/13 on recruitment of these cell populations to gills. This is unanticipated from mammals, where basophils and eosinophils infiltrate tissues coordinately (56).

The increase in the fraction of M1 macrophages as well as enhanced levels of *tnfα* and *il6* transcripts in the gills of *il4/13a;b^−/−^* double mutants indicate an inflammatory phenotype. Recent studies have identified subsets of polarized macrophages in zebrafish, describing a pro-inflammatory M1-like phenotype and anti-inflammatory M2-like phenotype, resembling the situation found in mammals (57). Our data suggest that IL-4/13A and IL-4/13B have a conserved function with mammalian IL-4 and IL-13 in driving macrophage polarization, as found in other teleost fish (58–61). The increase of IFN-γ in the gills of *il4/13a;b^−/−^* double mutants might also contribute to the induction of M1 macrophages, as previously demonstrated (62) and to the inflammatory phenotype. All together these findings indicate that loss of *il4/13a* and *il4/13b* leads to an inflammatory phenotype in the gills. This agrees with the phenotype observed in larvae, further supporting an anti-inflammatory function for zebrafish *il4/13* paralogues.

Despite induction of proinflammatory signals, loss of *il4/13a* and *il4/13b* did not affect tissue morphology, suggesting that there might be further regulatory mechanisms in place to avoid disruption of homeostasis in the gills. IL-10 is known to control mammalian immune responses in order to avoid inflammation and maintain tissue homeostasis. We examined the gills of *il10*^*e46/e46*^ fish and found that IL-10 might be fundamental in maintaining homeostasis in the gills. Transcriptome analysis revealed evidence of smoldering inflammation in the gills of *il10*^*e46/e46*^ fish. In mammals, lack of IL-10 leads to the development of spontaneous inflammation in the skin, intestine and lungs (63). In this context, zebrafish IL-10 might have a similar function to its mammalian counterpart.

Recent work has described the use of Resiquimod (R848) in inducing inflammation and cytokine response in zebrafish gills (33), highlighting similarities with mouse and human nasal mucosa. We used Resiquimod to induce inflammation in the gills and found that loss of both *il4/13a* and *il4/13b* lead to an enhanced response in the gills of double mutants, which appeared to be type-1-driven, in agreement with transcriptome data in the steady-state. In mammals, IL-4 and IL-13 act together to ensure a successful Th2 inflammatory response, and a Th1 phenotype with increased expression of IFN-γ is observed in IL-4/IL-13 deficient mice (44). We infer from our data that in the absence of functional IL-4/13A and IL-4/13B a Th1-like response dominates in zebrafish, indicating a similar function of these cytokines to their mammalian counterparts.

IL-10 appeared to be fundamental in the maintenance of gill homeostasis and its loss enhanced the inflammatory response to Resiquimod in the gills, supporting a potent anti-inflammatory function for this cytokine. An anti-inflammatory role for IL-10 has been previously shown in carp (64) and in zebrafish gut (65). Moreover, suppression of Th1 cell response and cytokine production was reported in a zebrafish *M.marinum* model (38). For the first time to our knowledge, we addressed the function of zebrafish IL-10 in the gills and demonstrated its importance in preventing inflammation in this mucosal tissue.

Previous studies indicate that fish IL-4/13 paralogues might have divergent functions, with IL-4/13A providing a basal level of type-2 immunity and IL-4/13B required for specific T cell mediated immunity (16). Our findings indicate that zebrafish *il4/13* paralogues have a redundant function in suppressing inflammation and maintaining a Th2 phenotype in the gills. Further investigation would be needed to discover unique functions for IL-4/13 cytokines in zebrafish immunity and our RNA-seq data will be a useful resource in this respect. We propose that the *il4/13a^−/−^*, *il4/13b^−/−^* and *il4/13a;b^−/−^* zebrafish mutants would be valuable models to investigate the function of *il4/13* paralogues in the context of infection and disease. This would further our knowledge on fish immunology and provide comparative models to answer outstanding questions in mammals.

## Acknowledgments

We thank our colleagues Peter Walker and Grace Bako (Histology Facility), Dr Andy Hayes (Genomic Technologies), Peter March (Bioimaging) and the staff from the Biological Service Unit (BSU) (all Faculty of Biology, Medicine and Health, The University of Manchester) for their technical assistance. We also thank the Central Biomedical Services within Imperial College London for their assistance. We thank Simon Johnston and Stephen Renshaw (University of Sheffield) for providing *il10*^*e46/e46*^ mutant zebrafish and *mpx* RNA probe, respectively. The study was begun using funding from Biotechnology and Biological Sciences Research Council Grant BB/L007401/1 to A.H. Federica Bottiglione was funded by a studentship from the Wellcome Trust (Grant 102171/Z/13/Z).

## References

1. Junttila IS. Tuning the cytokine responses: An update on interleukin (IL)-4 and IL-13 receptor complexes. Front Immunol. 2018;9:888.

2. Chen F, Liu Z, Wu W, Rozo C, Bowdridge S, Rooijen N Van, Urban JF Jr, Wynn TA and Gause WC. An essential role for Th2-type responses in limiting acute tissue damage during helminth infection. 2012;18(2):260–6.

3. Neill DR, Wong SH, Bellosi A, Flynn RJ, Daly M, Langford TK, Bucks C, Kane CM, Fallon PG, Pannell R, Jolin HE, McKenzie AN. Nuocytes represent a new innate effector leukocyte that mediates type-2 immunity. Nature. 2010;464(7293):1367–70.

4. McCormick SM, Heller NM. Commentary: IL-4 and IL-13 receptors and signaling. Cytokine 2015;75(1):38–50.

5. Gordon S. Alternative activation of macrophages. Nat Rev Immunol 2003;3(1):23–35.

6. Kara EE, Comerford I, Fenix KA, Bastow CR, Gregor CE, McKenzie DR, McColl SR. Tailored Immune Responses: Novel Effector Helper T Cell Subsets in Protective Immunity. PLoS Pathog. 2014;10(2).

7. Prete G Del. Human Th1 and Th2 lymphocytes: their role in the pathophysiology of atopy. Allergy. 1992;47(5):450–5.

8. Cope A, Le Friec G, Cardone J, Kemper C. The Th1 life cycle: Molecular control of IFN-γ to IL-10 switching. Trends Immunol. 2011;32(6):278–86.

9. Saraiva M, O’Garra A. The regulation of IL-10 production by immune cells. Nat Rev Immunol. 2010;10(3):170–81.

10. Barnes MJ, Powrie F. Regulatory T Cells Reinforce Intestinal Homeostasis. Immunity. 2009;31(3):401–11.

11. Yamaguchi T, Takizawa F, Fischer U, Dijkstra J. Along the Axis between Type 1 and Type 2 Immunity; Principles Conserved in Evolution from Fish to Mammals. Biology (Basel) 2015;4(4):814–59.

12. Venkatesh B, Lee AP, Ravi V, Maurya AK, Lian MM, Swann JB, Ohta Y, Flajnik MF, Sutoh Y, Kasahara M, Hoon S, Gangu V, Roy SW, Irimia M, Korzh V, Kondrychyn I, Lim ZW, Tay B-H, Tohari S, Kong KW, Ho S, Lorente-Galdos B, Quilez J, Marques-Bonet T, Raney BJ, Ingham PW, Tay A, Hillier LW, Minx P, Boehm T, Wilson RK, Brenner S, Warren WC. Elephant shark genome provides unique insights into gnathostome evolution. Nature. 2014;505(7482):174–9.

13. Gomez D, Sunyer J, Salinas I. The mucosal immune system of fish. Fish Shellfish Immunol. 2008;35(6):1729–39.

14. Campos-Perez JJ, Ward M, Grabowski PS, Ellis AE, Secombes CJ. The gills are an important site of iNOS expression in rainbow trout Oncorhynchus mykiss after challenge with the Gram-positive pathogen Renibacterium salmoninarum. Immunology. 2000;99(1):153–61.

15. Salinas I, Zhang YA, Sunyer JO. Mucosal immunoglobulins and B cells of teleost fish. Dev Comp Immunol. 2011;35(12):1346–65.

16. Wang T, Johansson P, Abós B, Holt A, Tafalla C, Jiang Y, Wang A, Xu Q, Qi Z, Huang W, Costa MM, Diaz-Rosales P, Holland JW, Secombes CJ. First in-depth analysis of the novel Th2-type cytokines in salmonid fish reveals distinct patterns of expression and modulation but overlapping bioactivities. Oncotarget. 2016;7(10):10917–46.

17. Mitra S, Alnabulsi A, Secombes CJ, Bird S. Identification and characterization of the transcription factors involved in T-cell development, t-bet, stat6 and foxp3, within the zebrafish, Danio rerio. FEBS J. 2010;277(1):128–47.

18. Ohtani M, Hayashi N, Hashimoto K, Nakanishi T, Dijkstra JM. Comprehensive clarification of two paralogous interleukin 4/13 loci in teleost fish. Immunogenetics. 2008;60(7):383–97.

19. Avery S, Rothwell L, Degen WDJ, Schijns VE, Young J, Kaufman J, Kaiser P. Characterization of the first nonmammalian T2 cytokine gene cluster: The cluster contains functional single-copy genes for IL-3, IL-4, IL-13, and GM-CSF, a gene for IL-5 that appears to be a pseudogene, and a gene encoding another cytokinelike transcript, KK34. J Interf Cytokine Res. 2004;24(10):600–10.

20. Takizawa F, Koppang EO, Ohtani M, Nakanishi T, Hashimoto K, Fischer U, Dijkstra JM. Constitutive high expression of interleukin-4/13A and GATA-3 in gill and skin of salmonid fishes suggests that these tissues form Th2-skewed immune environments. Mol Immunol. 2011;48(12-13):1360–8.

21. Dee CT, Nagaraju RT, Athanasiadis EI, Gray C, Fernandez del Ama L, Johnston SA, Secombes CJ, Cvejic A, Hurlstone AF. CD4-Transgenic Zebrafish Reveal Tissue-Resident Th2- and Regulatory T Cell-like Populations and Diverse Mononuclear Phagocytes. J Immunol. 2016.

22. Zou J, Clark MS, Secombes CJ. Characterisation, expression and promoter analysis of an interleukin 10 homologue in the puffer fish, Fugu rubripes. Immunogenetics. 2003;55(5):325–35.

23. Savan R, Igawa D, Sakai M. Cloning, characterization and expression analysis of interleukin-10 from the common carp, Cyprinus carpio L. Eur J Biochem. 2003;270(23):4647–54.

24. Inoue Y, Kamota S, Ito K, Yoshiura Y, Ototake M, Moritomo T, Nakanishi T. Molecular cloning and expression analysis of rainbow trout (Oncorhynchus mykiss) interleukin-10 cDNAs. Fish Shellfish Immunol. 2005;18(4):335–44.

25. Zhang DC, Shao YQ, Huang YQ, Jiang SG. Cloning, characterization and expression analysis of interleukin-10 from the zebrafish (Danio rerio). J Biochem Mol Biol. 2005;38(5):571–6.

26. Kettleborough RNW, Busch-nentwich EM, Harvey SA, Dooley CM, Bruijn E De, Eeden F Van, Sealy I, White RJ, Herd C, Nijman IJ, Fènyes F, Mehroke S, Scahill C, Gibbons R, Wali N, Carruthers S, Hall A, Yen J, Cuppen E, Stemple DL. A systematic genome-wide analysis of zebrafish protein-coding gene function. Nature 2013;496(7446):494–7.

27. Thisse C, Thisse B. High-resolution in situ hybridization to whole-mount zebrafish embryos. Nat Protoc. 2008;3(1):59–69.

28. Bolger AM, Lohse M, Usadel B. Trimmomatic: A flexible trimmer for Illumina sequence data. Bioinformatics. 2014;30(15):2114–20.

29. Dobin A, Davis CA, Schlesinger F, Drenkow J, Zaleski C, Jha S, Batut P, Chaisson M, Gingeras TR. STAR: Ultrafast universal RNA-seq aligner. Bioinformatics. 2013;29(1):15–21.

30. Love MI, Huber W, Anders S. Moderated estimation of fold change and dispersion for RNA-seq data with DESeq2. Genome Biol. 2014;15(12):1–21.

31. Kuleshov MV, Jones MR, Rouillard AD, Fernandez NF, Duan Q, Wang Z, Koplev S, Jenkins SL, Jagodnik KM, Lachmann A, McDermott MG, Monteiro CD, Gundersen GW, Ma’ayan A. Enrichr: a comprehensive gene set enrichment analysis web server 2016 update. Nucleic Acids Res. 2016;44(W1):W90–7.

32. Subramanian A, Tamayo P, Mootha VK, Mukherjee S, Ebert BL, Gillette MA, Paulovich A, Pomeroy SL, Golub TR, Lander ES, Mesirov JP. Gene set enrichment analysis: a knowledge-based approach for interpreting genome-wide expression profiles. Proc Natl Acad Sci U S A. 2005;102(43):15545–50.

33. Progatzky F, Jha A, Wane M, Thwaites RS, Makris S, Shattock RJ, Johansson C, Openshaw PJ, Bugeon L, Hansel TT, Dallman MJ. Induction of innate cytokine responses by respiratory mucosal challenge with R848 in zebrafish, mice, and humans. J Allergy Clin Immunol. 2019;144(1):342–345.e7.

34. Nagy E, Maquat LE. A rule for termination-codon position within intron-containing genes: When nonsense affects RNA abundance. Trends Biochem Sci. 1998;23(6):198–9.

35. El-Brolosy MA, Kontarakis Z, Rossi A, Kuenne C, Günther S, Fukuda N, Kikhi K, Boezio GLM, Takacs CM, Lai SL, Fukuda R, Gerri C, Giraldez AJ, Stainier DYR. Genetic compensation triggered by mutant mRNA degradation. Nature. 2019;568(7751):193–7.

36. Stocchi V, Wang T, Randelli E, Mazzini M, Gerdol M, Pallavicini A, Secombes CJ, Scapigliati G, Buonocore F. Evolution of Th2 responses: Characterization of IL-4/13 in sea bass (Dicentrarchus labrax L.) and studies of expression and biological activity. Sci Rep. 2017;7(1):1–15.

37. Herbomel P, Thisse B, Thisse C. Zebrafish early macrophages colonize cephalic mesenchyme and developing brain, retina, and epidermis through a M-CSF receptor-dependent invasive process. Dev Biol. 2001;238(2):274–88.

38. Harjula SKE, Ojanen MJT, Taavitsainen S, Nykter M, Rämet M. Interleukin 10 mutant zebrafish have an enhanced interferon gamma response and improved survival against a Mycobacterium marinum infection. Sci Rep. 2018;8(1):1–17.

39. Hemmi H, Kaisho T, Takeuchi O, Sato S, Sanjo H, Hoshino K, Horiuchi T, Tomizawa H, Takeda K, Akira S. Small-antiviral compounds activate immune cells via the TLR7 MyD88-dependent signaling pathway. Nat Immunol. 2002;3(2):196–200.

40. Gomez D, Sunyer JO, Salinas I. The mucosal immune system of fish: The evolution of tolerating commensals while fighting pathogens. Fish Shellfish Immunol. 2013;35(6):1729–39.

41. Wang T, Secombes CJ. The cytokine networks of adaptive immunity in fish. Fish Shellfish Immunol. 2013;35(6):1703–18.

42. Shinn AP, Pratoomyot J, Bron JE, Paladini G, Brooker EE, Brooker AJ. Economic costs of protistan and metazoan parasites to global mariculture. Parasitology. 2015;142(1):196–270.

43. Herrero A, Thompson KD, Ashby A, Rodger HD, Dagleish MP. Complex Gill Disease: an Emerging Syndrome in Farmed Atlantic Salmon (Salmo salar L.). J Comp Pathol 2018;163:23–8.

44. Mckenzie BGJ, Fallon PG, Emson CL, Grencis RK, Mckenzie AN. Simultaneous disruption of interleukin (IL)-4 and IL-13 defines individual roles in T helper cell type 2 – mediated Responses. J Exp Med 1999;189(10):0–7.

45. Novoa B, Figueras A. Zebrafish: Model for the study of inflammation and the innate immune response to infectious diseases. Vol. 946, Advances in Experimental Medicine and Biology. 2012. 253–275 p.

46. Girard D, Paquin R, Beaulieu AD. Responsiveness of human neutrophils to interleukin-4: Induction of cytoskeletal rearrangements, de novo protein synthesis and delay of apoptosis. Biochem J. 1997;325(1):147–53.

47. Girard D, Ratthé C, Ennaciri J, Garcs Gonalves DM, Chiasson S. Interleukin (IL)-4 induces leukocyte infiltration in vivo by an indirect mechanism. Mediators Inflamm. 2009;2009:1–11.

48. Egholm C, Heeb LEM, Impellizzieri D, Boyman O. The regulatory effects of interleukin-4 receptor signaling on neutrophils in type 2 immune responses. Front Immunol. 2019;10(OCT):1–14.

49. Heeb LEM, Egholm C, Impellizzieri D, Ridder F, Boyman O. Regulation of neutrophils in type 2 immune responses. Curr Opin Immunol. 2018;54:115–22.

50. Heeb LEM, Egholm C, Boyman O. Evolution and function of interleukin-4 receptor signaling in adaptive immunity and neutrophils. Genes Immun 2020. 2020;1–7.

51. Zhu L, Pan P, Fang W, Shao J, Xiang L. Essential Role of IL-4 and IL-4Rα Interaction in Adaptive Immunity of Zebrafish: Insight into the Origin of Th2- like Regulatory Mechanism in Ancient Vertebrates. J Immunol. 2012;188(11):5571–84.

52. Marcos-López M, Calduch-Giner JA, Mirimin L, MacCarthy E, Rodger HD, O’Connor I, Sitjà-Bobadilla A, Pèrez-Sànchez J, Piazzon MC. Gene expression analysis of Atlantic salmon gills reveals mucin 5 and interleukin 4/13 as key molecules during amoebic gill disease. Sci Rep. 2018;8(1):1–15.

53. Voehringer D, Reese TA, Huang X, Shinkai K, Locksley RM. Type 2 immunity is controlled by IL-4/IL-13 expression in hematopoietic non-eosinophil cells of the innate immune system. J Exp Med. 2006;203(6):1435–46.

54. Odaka T, Suetake H, Maeda T, Miyadai T. Teleost Basophils Have IgM-Dependent and Dual Ig-Independent Degranulation Systems. J Immunol. 2018;200(8):2767–76.

55. Balla KM, Lugo-Villarino G, Spitsbergen JM, Stachura DL, Hu Y, Bañuelos K, Romo-Fewell O, Aroian RV, Traver D. Eosinophils in the zebrafish: Prospective isolation, characterization, and eosinophilia induction by helminth determinants. Blood. 2010;116(19):3944–54.

56. Voehringer D, Shinkai K, Locksley RM. Type 2 immunity reflects orchestrated recruitment of cells committed to IL-4 production. Immunity. 2004;20(3):267–77.

57. Nguyen-Chi M, Laplace-Builhe B, Travnickova J, Luz-Crawford P, Tejedor G, Phan QT, Duroux-Richard I, Levraud JP, Kissa K, Lutfalla G, Jorgensen C, Djouad F. Identification of polarized macrophage subsets in zebrafish. Elife. 2015;4:e07288.

58. Jiang X, Wang J, Wan S, Xue Y, Sun Z, Cheng X, Gao Q, Zou J. Distinct expression profiles and overlapping functions of IL-4/13A and IL-4/13B in grass carp (Ctenopharyngodon idella). Aquac Fish. 2020;5(2):72–9.

59. Wiegertjes GF, Wentzel AS, Spaink HP, Elks PM, Fink IR. Polarization of immune responses in fish: The ‘macrophages first’ point of view. Mol Immunol. 2016;69:146–56.

60. Hodgkinson JW, Fibke C, Belosevic M. Recombinant IL-4/13A and IL-4/13B induce arginase activity and down-regulate nitric oxide response of primary goldfish (Carassius auratus L.) macrophages. Dev Comp Immunol. 2017;67:377–84.

61. Joerink M, Ribeiro CMS, Stet RJM, Hermsen T, Savelkoul HFJ, Wiegertjes GF. Head Kidney-Derived Macrophages of Common Carp (Cyprinus carpio L.) Show Plasticity and Functional Polarization upon Differential Stimulation. J Immunol. 2006;177(1):61–9.

62. Zou J, Carrington A, Collet B, Dijkstra JM, Yoshiura Y, Bols N, Secombes C. Identification and Bioactivities of IFN-γ in Rainbow Trout Oncorhynchus mykiss: the first Th1-type cytokine characterized functionally in fish. J Immunol. 2005;175(4):2484–94.

63. Rubtsov YP, Rasmussen JP, Chi EY, Fontenot J, Castelli L, Ye X, Treuting P, Siewe L, Roers A, Henderson WR Jr, Muller W, Rudensky AY. Regulatory T cell-derived interleukin-10 limits inflammation at environmental interfaces. Immunity. 2008;28(4):546–58.

64. Piazzon MC, Savelkoul HSJ, Pietretti D, Wiegertjes GF, Forlenza M. Carp Il10 has anti-inflammatory activities on phagocytes, promotes proliferation of memory T cells, and regulates B cell differentiation and antibody secretion. J Immunol. 2015;194(1):187–99.

65. Coronado M, Solis CJ, Hernandez PP, Feijóo CG. Soybean meal-induced intestinal inflammation in zebrafish is T cell-dependent and has a Th17 cytokine profile. Front Immunol. 2019;10:1–13.

