## Supplemental Figure for "Zebrafish IL-4-like cytokines and IL-10 suppress inflammation but only IL-10 is essential for gill homeostasis"

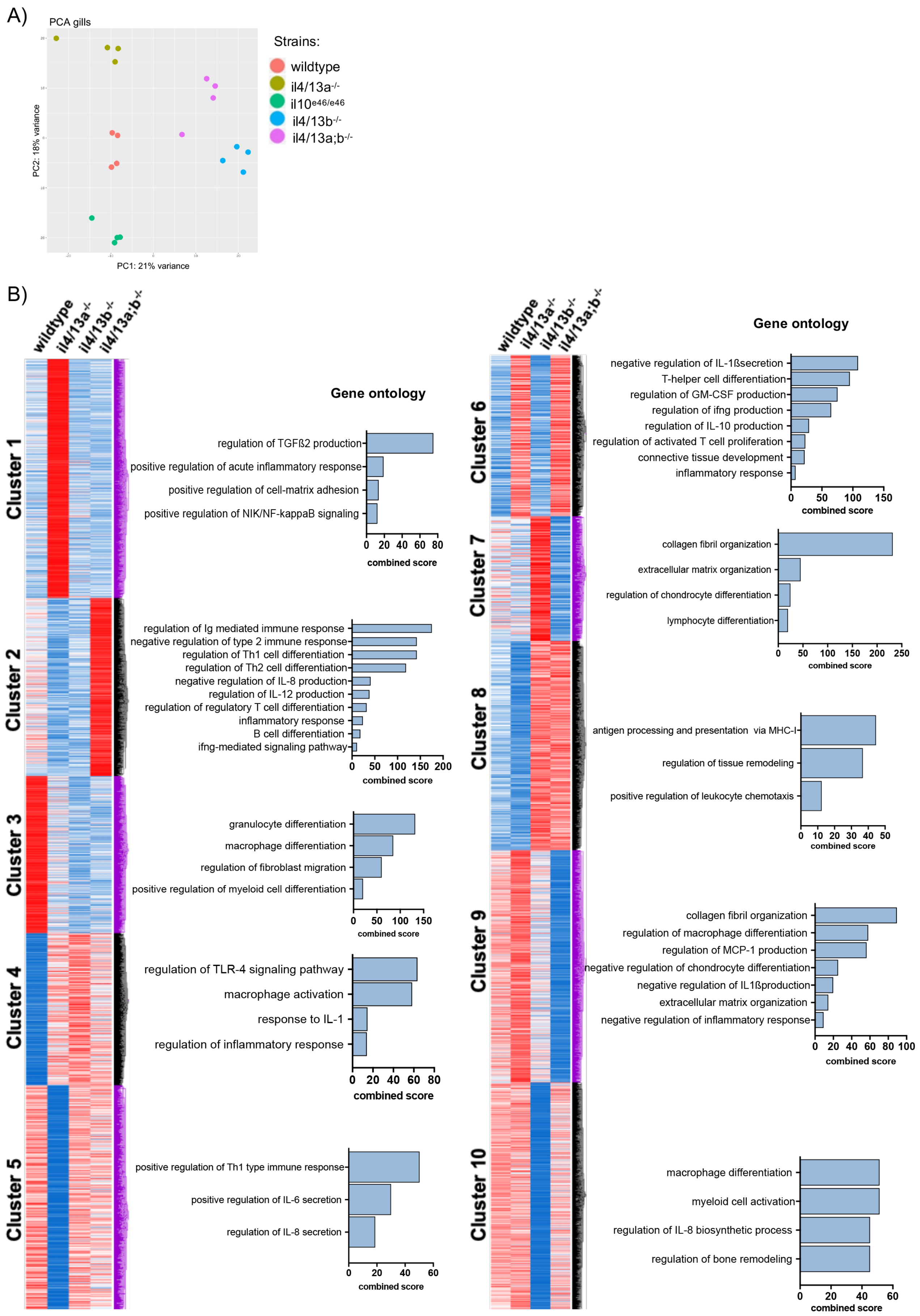
A)
PCA gills
PC2: 18% variance
PC1: 21% variance
Strains:
wildtype
il4/13a-/-
il10e46/e46
il4/13b-/-
il4/13a;b-/-
B)
wildtype
il4/13a-/-
il4/13b-/-
il4/13a;b-/-
Cluster 1
Cluster 2
Cluster 3
Cluster 4
Cluster 5
Cluster 6
Cluster 7
Cluster 8
Cluster 9
Cluster 10
Gene ontology
regulation of TGFß2 production
positive regulation of acute inflammatory response
positive regulation of cell-matrix adhesion
positive regulation of NIK/NF-kappaB signaling
combined score
regulation of Ig mediated immune response
negative regulation of type 2 immune response
regulation of Th1 cell differentiation
regulation of Th2 cell differentiation
negative regulation of IL-8 production
regulation of IL-12 production
regulation of regulatory T cell differentiation
inflammatory response
B cell differentiation
ifng-mediated signaling pathway
granulocyte differentiation
macrophage differentiation
regulation of fibroblast migration
positive regulation of myeloid cell differentiation
regulation of TLR-4 signaling pathway
macrophage activation
response to IL-1
regulation of inflammatory response
positive regulation of Th1 type immune response
positive regulation of IL-6 secretion
regulation of IL-8 secretion
negative regulation of IL-1ßsecretion
T-helper cell differentiation
regulation of GM-CSF production
regulation of ifng production
regulation of IL-10 production
regulation of activated T cell proliferation
connective tissue development
inflammatory response
collagen fibril organization
extracellular matrix organization
regulation of chondrocyte differentiation
lymphocyte differentiation
antigen processing and presentation via MHC-I
regulation of tissue remodeling
positive regulation of leukocyte chemotaxis
collagen fibril organization
regulation of macrophage differentiation
regulation of MCP-1 production
negative regulation of chondrocyte differentiation
negative regulation of IL1ßproduction
extracellular matrix organization
negative regulation of inflammatory response
macrophage differentiation
myeloid cell activation
regulation of IL-8 biosynthetic process
regulation of bone remodeling

**Supplemental Figure 1** (A) Principal components analysis (PCA) plots representing the distribution of gills samples. PCA plots were generated using DESeq2_1.16.1. (B) Clustering of differentially expressed genes identified in gills harvested from 6 month old fish. Blue indicates low expression, red indicates high expression and white indicates unchanged expression. Bar graphs show significantly enriched ontologies for each cluster of genes.
